# Molossus molossus bats encompass at least two distinct genera of totivirus-like

**DOI:** 10.1101/2023.07.07.548100

**Authors:** Roseane da Silva Couto, Endrya do Socorro Foro Ramos, Wandercleyson Uchôa Abreu, Luis Reginaldo Ribeiro Rodrigues, Luis Fernando Marinho, Vanessa dos Santos Morais, Fabiola Villanova, Ramendra Pati Pandey, Xutao Deng, Eric Delwart, Antonio Charlys da Costa, Elcio Leal

**Author notes:** Correspondence (E.L.). These authors contributed equally to this work (Roseane da Silva Couto, Endrya do Socorro Foro Ramos, Wandercleyson Uchôa Abreu). These authors jointly supervised this work (Antonio Charlys da Costa, Elcio Leal). (R.S.C.); (E.d.S.F.R.); (F.V.). (W.U.A.). (L.R.R.R.). (L.F.M.). (V.d.S.M.); (A.C.d.C.). (R.P.P.). (E.D.) (X.D.).

## Abstract

The Totiviridae family of viruses has a unique genome consisting of double-stranded RNA with two open reading frames that encode the capsid protein (Cap) and the RNA-dependent RNA polymerase (RdRp). Most virions in this family are isometric in shape, approximately 40 nm in diameter, and lack envelope. There are five genera within this family, including Totivirus, *Victorivirus, Giardiavirus, Leishmaniavirus*, and *Trichomonasvirus*. While *Totivirus* and *Victorivirus* primarily infect fungi, *Giardiavirus, Leishmaniavirus*, and *Trichomonasvirus* infect diverse hosts, including protists, insects, and vertebrates. Recently, new totivirus-like species have been discovered in fish and plant hosts, and through metagenomic analysis, a novel totivirus-like virus (named Tianjin totivirus) has been isolated from bat guano. Interestingly, Tianjin totivirus causes cytopathic effects in insect cells but cannot grow in mammalian cells, suggesting that it infects insects consumed by insectivorous bats. In this study, we used next-generation sequencing and identified totivirus-like viruses in liver tissue from *Molossus molossus* bats () in the Amazon region of Brazil. Comparative phylogenetic analysis based on the RNA-dependent RNA polymerase region revealed that the viruses identified in Molossus bats belong to two distinct phylogenetic clades, possibly comprising different genera within the Totiviridae family. Notably, the mean similarity between Tianjin totivirus and the totiviruses identified in Molossus bats is less than 18%. These findings suggest that the diversity of totiviruses in bats is more extensive than previously recognized and highlight the potential for bats to serve as reservoirs for novel toti-like viruses.

## Introduction

Bats are unique animals due to their extensive viral diversity, which distinguishes them from other species [1–6]. They have long been associated with many viral families and genera, such as *Paramyxoviridae*, Filoviridae, Lyssavirus, and *Henipavirus* [7–17]. Their ability to fly long distances and their diverse feeding habits make it easier for them to acquire and spread viruses across remote areas and to transmit them to other species. Additionally, their social structures and behaviors contribute to virus transmission and persistence within bat populations [4,18– 20]. However, changes in the environment, such as urbanization, agricultural intensification, and deforestation, have altered the composition and dynamics of bat communities [21]. Studies have shown an overall decline in species richness and relative abundance associated with urbanization [22]. Nevertheless, insectivorous bats tend to thrive in large urban environments [4,18,19]. Furthermore, the diversity of bat habitats can influence both microbe transmission and persistence in bat communities [2, 22–24].

Viral metagenomics research in bats has mainly focused on North American and Eurasian bat communities [21,25–27]. However, studies in the Amazon region have enabled the identification of numerous viruses using conventional techniques or high-throughput sequencing [7–9,9,12,14,14,16,28,29]. For instance, a metagenomic study conducted in French Guiana found various RNA viruses in fecal samples of Molussus bats (also known as velvety free-tailed bat or Pallas’s mastiff bat), including some short sequences of viruses belonging to the Totiviridae family [30]. Another study performed in carcasses of deceased bats in Germany, also found short sequence of totiviruses [31]. Researchers in China were able isolate one toti-like virus in insect cells from guano samples of the insectivorous Myotis bats [32].

Recent studies have shed light on the diverse range of insect viruses found in bat droppings through Next Generation Sequencing (NGS) [21,33]. NGS has facilitated the isolation and characterization of several new totivirus-like viruses that have yet to be classified by the International Committee on Taxonomy of Viruses (ICTV). The *Totiviridae* family is a diverse group of RNA viruses that infect both protozoa and fungi, with five identified genera to date. The family comprises 28 species, each of which contains a single molecule of double-stranded RNA (dsRNA) ranging in size from 4.6 to 7.0 kbp. The virus genome consists of two frames, with the 5’ ORF encoding the capsid protein (CP) and the 3’ ORF encoding the RNA-dependent RNA polymerase (RdRP) gene [34].

Viruses in the Totiviridae family are remarkably diverse in their ability to infect hosts ranging from fungi to protozoa [35–43]. The current recognized genera include Giardiavirus, *Leishmaniavirus, Totivirus, Victorivirus*, and *Trichomonasvirus* [35,44–49]. While those in the genera *Totivirus* and *Victorivirus* infect fungi, those in *Giardiavirus, Leishmaniavirus*, and *Trichomonasvirus* infect protozoa. Recently, new viruses have been identified in shrimp, fish, and mosquitoes that have similar genomic structures and morphology but low similarity to members of other genera in the Totiviridae family [47,50]. Researchers have also proposed three new genera within the Totiviridae family, namely *Artivirus, Pistolvirus*, and *Tricladiviris* [39,45,48,49,51,52]. For instance, Artivirus has been found to infect arthropods such as the Atlantic blue crab [53] and the mosquito Armigere subalbatus totivirus [54]. *Pistolvirus* has been found to infect fish species such as the Atlantic salmon and golden carp [39,52]. *Tricladiviris* has been found to infect planaria. Additionally, new toti-like viruses have been discovered in Eysarcoris guttigerus, with genomic characteristics similar to Sanya orius sauteri totivirus 2. Arboreal ants Camponotus yamaokai also carry a virus that is potentially a new totivirus, but it is not related to the Totiviruses identified in arthropods [55].

In our study, we utilized a metagenomic next-generation sequencing (NGS) approach to comprehensively examine viruses present in the insectivorous *Molossus molossus* bats that were captured in the urban areas of Santarém city, located in northern Brazil. Our analysis yielded forty-seven contigs, including three near-complete genomes, which showed an average amino acid identity of 40% with a closely related totivirus-like previously reported.

## Materials and Methods

### Sample collection

We captured a total of 47 bats in Caranazal (latitude 2°26’10”S and longitude 54°43’49”W), a municipality in the Santarém region of Pará state in the Lower Amazon Mesoregion. These bats were identified to *Molossus molossus* (Family *Molossidae*) based on external characters. To obtain the sequences, individual bats were euthanized for sample collection. We administered xylazine hydrochloride (1mg/kg) and ketamine hydrochloride (1-2mg/kg) via intramuscular injection to induce anaesthesia, followed by intracardiac phenobarbital (40mg/kg) once the animals had lost consciousness. The liver samples were then collected for further analysis.

To carry out this research, approval was obtained from the Animal Use Ethics Committee of the Federal University of Western Pará (CEUA/UFOPA) under number 0220220128 and from the Biodiversity Information and Authorization System (SISBIO -18313-1) for capturing Chiroptera. The necropsy was performed at the Animal Morphology Laboratory of the Federal University of Western Pará, following the institutional biosafety norms.

### Processing of samples

To extract viral particles from liver tissue, we prepared pools of organ pieces from five different animals. These pools were named F1, F2 and so on. The tissue was first macerated in a tissue disruptor and then diluted in 500 μL of Hanks Buffered Saline Solution (HBSS). Next, the sample was added to a 2 mL tube containing lysis matrix C (MP Biomedicals, Santa Ana, CA, USA) and homogenized using a Vortex mixer. After removing the large debris, the supernatants were filtered through 0.45 μM filters (Merck Millipore, Billerica, MA, USA) to eliminate eukaryotic and bacterial cell particles. The resulting filtrates were placed in 1.5 mL screw cap tubes and centrifuged at 32,000 rpm for 5 minutes using a Beckman Coulter Optima LE-80 ultracentrifuge with a Heraeus Maximum rotor. This step allowed the viral particles to sediment, while the supernatant was carefully removed. The pellet (which may not be visible) was resuspended in 250 μL of PBS, and the samples were then ready for nuclease enzyme treatment. The filtrates were treated with DNase (concentration, 20 U/ml; Ambion) and RNase A (concentration, 0.1 mg/ml; Fermentas) at 37°C for 30 min to digest unprotected particle nucleic acids. Phi29 (Φ29) polymerase enzyme was also used for DNA circular amplification.

### Nucleic acid extraction (DNA/RNA)

After sample preparation, viral nucleic acids were extracted using the QIAamp Viral RNA Mini Kit (QIAGEN GmbH, QIAGEN Strasse 1, 40724 Hilden, Germany), which purifies RNA and DNA, and the steps were followed according to the instructions of the manufacturer.

### Preparation of libraries for the illumina platform

Library preparation was performed using the Nextera XT DNA Sample Preparation Kit (Illumina Inc.), following the manufacturer’s guidelines. The Agilent 2100 Bioanalyzer system and the KAPA kit were used to perform library quantification. For sequencing, samples were pooled (5 organ samples per pool). After preparing the libraries, they were sequenced on the Illumina NovaSeq-6000 platform to provide 250 bp (base pairs) paired reads (Illumina).

### Bioinformatics Analysis

Raw reads obtained from Illumina sequencing were pre-processed, where: a) final matched sequence records were removed from both ends, b) low quality sequences (raw data generated from reads with less than 100 bp in length), and the adapter and primer sequences were cut using VecScreen based on BLAST (Basic Local Alignment Search Tool) at the default parameters. Bioinformatics data were analyzed according to the previously described protocol [56]. The resulting contigs were compared using BLASTx and B LASTn to look for similarity to viral proteins and nucleotides, respectively, from the GenBank genetic sequence database (http://www.ncbi.nlm.nih.gov, accessed January 9, 2023). The best results of searches in BLAST (with the highest percentage of sequence similarity ever deposited in Genbank) were selected and, to reduce the number of random matches, E values (e-value) were defined in each search. Based on the bioinformatics pipeline used, no reads related to human, plant, fungal, or bacterial sequences were obtained. All sequences generated in this study were deposited in the Genbank with the accession numbers OR069303-OR069349.

### Genome annotation

Comparison of RNA virus sequences in studies was obtained from liver samples of M. molossus andsubmitted to the online database DIAMOND (Double Index Alignment of Next Generation Sequencing Data), in which homologous sequences are compared and the taxonomic classification of viruses is refined [57]. As well as the comparative analysis of the predicted gene sequences via the BLASTx online program, as it is a protein aligner using a DNA alignment. Therefore, the high similarity between query strings is described by an alignment. Based on the best result, the sequences were selected to be further aligned.

All sequences generated in this study were similar to reference sequences from Genbank classified in the Totiviridae family. Subsequently, these reference sequences were selected and aligned using the MAFFT software, v7. Complete or almost complete genomes were aligned. Alignments and edits were made using Ugene tool kit version 4.6 [58]. The RdRpol domains and motifs were predicted using InterProScan (https://www.ebi.ac.uk/interpro/search/sequence, accessed 10 January 2023) and Motif Finder (https://www.genome.jp/tools/motif, accessed January 12, 2023), respectively.

### Genetic distances

The genetic distance and its standard error were calculated using the composite model of maximum likelihood plus gamma correction and bootstrapping with 1000 repetitions. Distances were calculated using the MEGA X software [59]. To estimate the similarity of the sequences, a paired method was used, implemented in the SDT program [60]. The estimation of the similarity alignments of each unique pair of sequences was done using algorithms implemented in MUSCLE [61]. After calculating the identity score for each pair of sequences, the program uses the NEIGHBOR component of PHYLIP to calculate a tree. The rooted neighbor-joining phylogenetic tree orders of all sequences according to their probable degrees of evolutionary relatedness. The results are presented in a frequency distribution of paired identities in a graphical interface.

### Phylogenetic analysis

Phylogenetic trees were constructed using the maximum likelihood approach, and branching support was estimated using a bootstrap test with 1000 repetitions using the IQ-Tree tool [62]. The maximum clade credibility tree was obtained using a Bayesian coalescent approach, implemented in the Beast v1.10.4 software (https://github.com/beast-dev/beast-mcmc). We assumed a fixed clock and a constant rate of population size, and assumed that the evolutionary rate of a given site in a gene-sequence alignment was constant during evolution (homotachy). The WAG+I+G evolutionary model were used with the number of gamma categories set to 4. The runs were initiated using a random starting tree and a chain length of 100,000,000, with echo state to screen every 10,000 iterations, log parameters every 10,000 iterations, and a burn-in of 10%. To ensure convergence and an effective sample size (ESS) greater than 200, we examined the results using Tracer v1.7.1 (http://tree.bio.ed.ac.uk/software/tracer).

## Results

### Contigs quality and BlastX similarities

We successfully retrieved forty-seven contigs from three different pools (F1, F3, and F6), which were identified as totiviruses based on blast searches. The contigs’ quality, sequence size, and blast similarities are summarized in Table 1. The general quality of contigs was excellent due to the high coverage and the inclusion of a large number of reads in the assembly of each contig. BlastX comparative analysis showed that contigs identified in liver samples of Molossus bats had low amino acid identity in the NCBI database (mean 48%), with coverage ranging from 15% to 97% with theirs best-hit reference sequence. Three of the contigs (F1_001 F1_002 and F1_003) represent near full length genome of totiviruses.

**Table 1.**
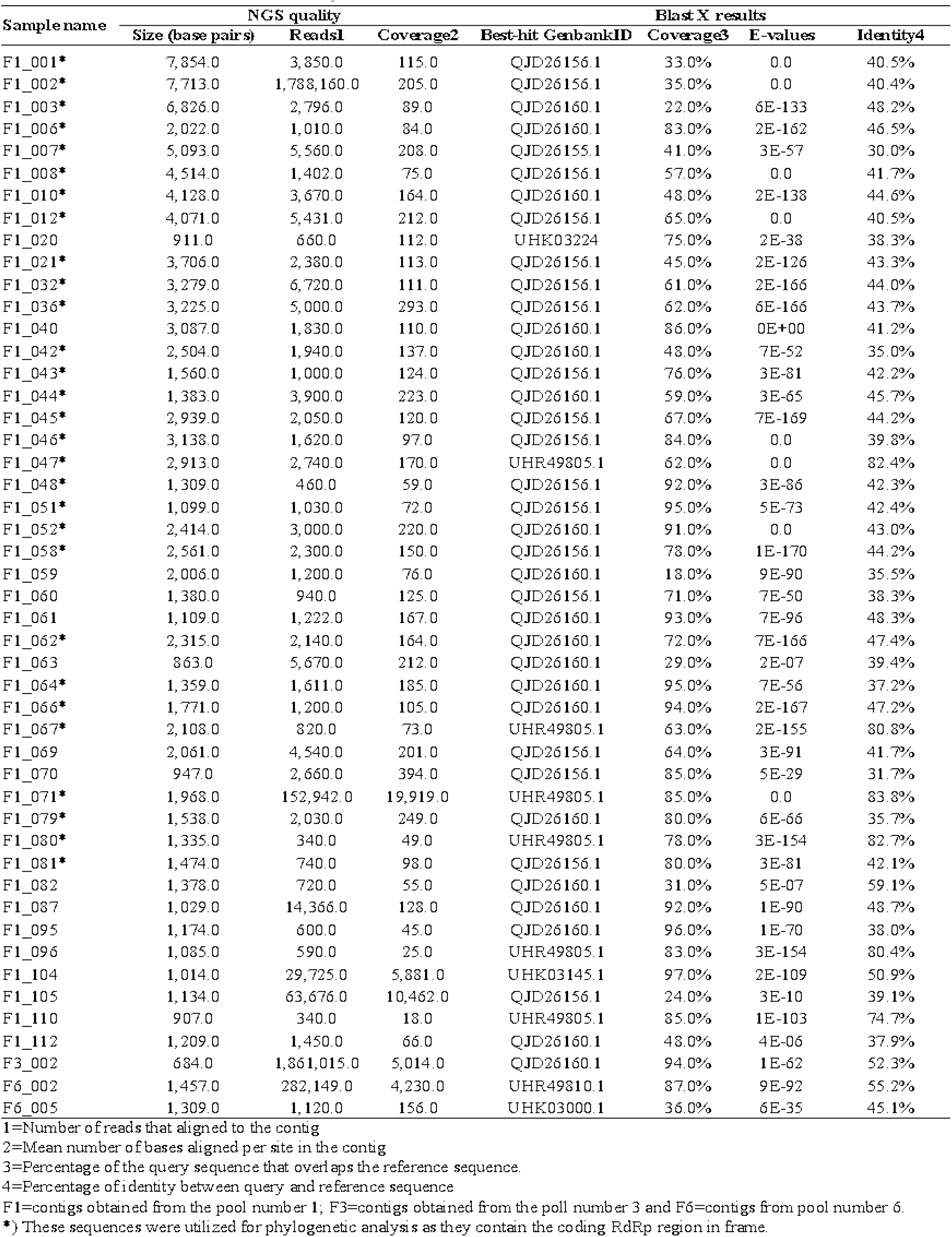
Characteristics of contigs identified Molossus bats.

### Genome annotation of totiviruses identified in bats

We have discovered three nearly complete genomes of totiviruses in liver samples collected from Molossus bats, specifically F1_001, F1_002 and F1_003. Using a BlastX search, we found that these two sequences share a degree of amino acid similarity with totiviruses previously identified in the helminth *Schistocephalus solidus*. To further explore this similarity, we compared the genome annotations of all these sequences (Fig 1). It is worth noting that the reference sequences MN803435 and MN803437, identified in *Schistocephalus solidus*, are related, but they have distinct genome maps. MN803437 has an additional open reading frame (orf), and the proteins have different sizes. For instance, the polymerase in MN803437 has 840 amino acids, while in MN803435, it has 953 amino acids. Both F1_001 and F1_003 contain three open reading frames (ORFs), with F1_003 having a notably short hypothetical polymerase sequence (only 486 amino acids). An important characteristic of these sequences is the presence of long non-coding intergenic regions, such as the 538bp region found between the second complete genome F1_002 has two hypothetical proteins, the first has 1456 residues and the second (probably RdRpol) has 1042 residues.

**Fig 1.**
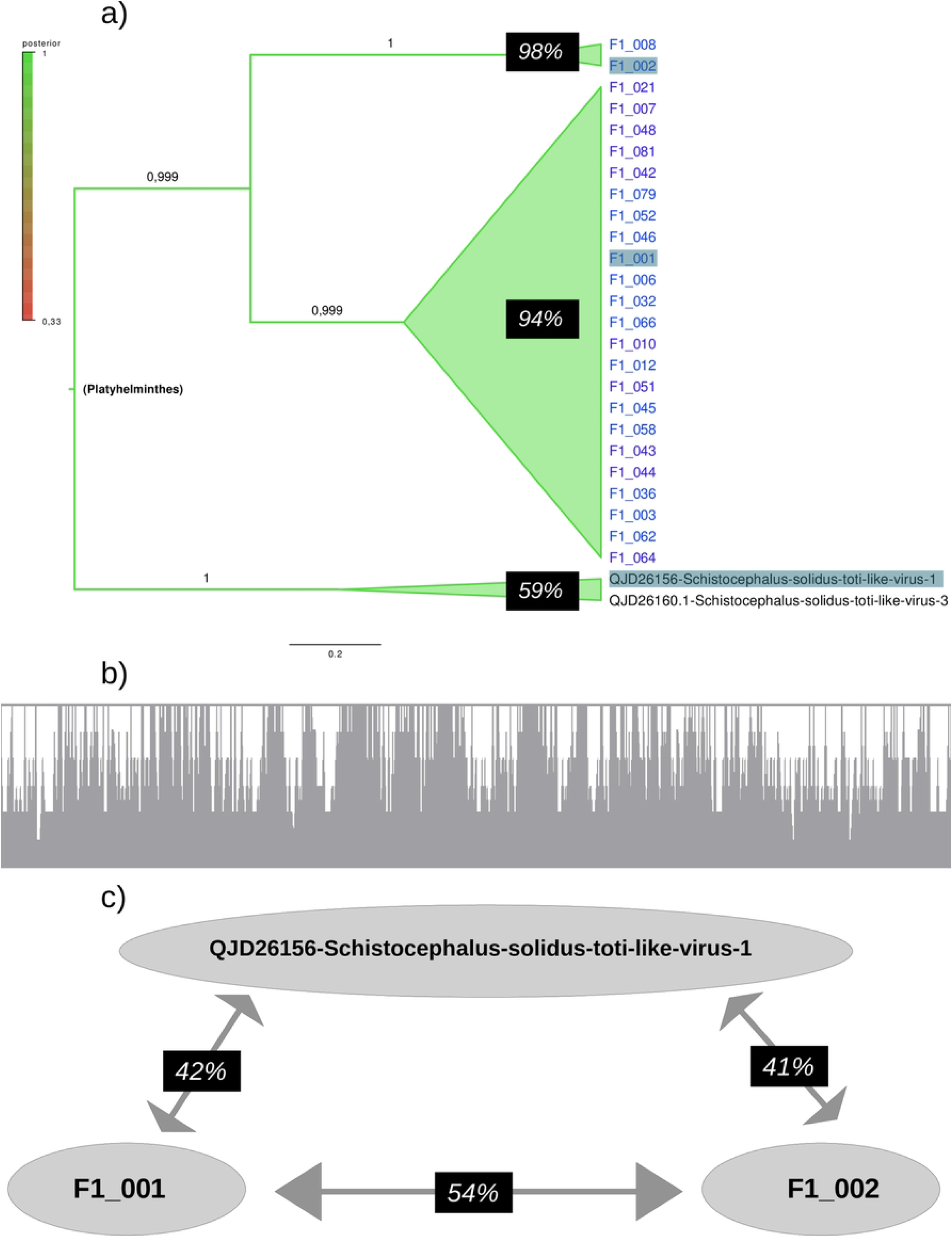
Genome map of totiviruses. Horizontal diagrams represent the location of hypothetical open reading frames (orf) in the genome of totiviruses. Upper panels indicate the orfs (cyan bars) detected in the genomes of sequences MN803435 and MN803437, identified in *Schistocephalus solidus*. Lower panels show the orfs (orange bars) detected in the genomes of F1_001, F1_002 and F1_003 identified in Molossus bats.

### RNA-dependent RNA-polymerase of totiviruses

Due to the limited similarities between our contigs and the reference sequences, we opted to analyze the motifs of the RNA-dependent RNA-polymerase region of totiviruses. To achieve this, we utilized the cognate viruses identified through the Blast search, as well as additional contigs found in Molossus bats (Fig 2). To illustrate our findings, we have included the motifs of RdRpol from three contigs that belong to different groups of totiviruses (as determined by our phylogenetic analysis). In addition to these motifs, we have also shown the motifs of their respective best-hits. It is important to note that even though the RdRpol sequences identified in this study are highly divergent (more than 50%), all of the motifs of the polymerase are present. Typically, cognate sequences share the same pattern, with one exception being the sequence F1_001, which possesses an additional motif (indicated by an arrow in Figure 2a) when compared to its best-hit reference QJD26160, identified in *Schistocephalus solidus*.

**Fig 2.**
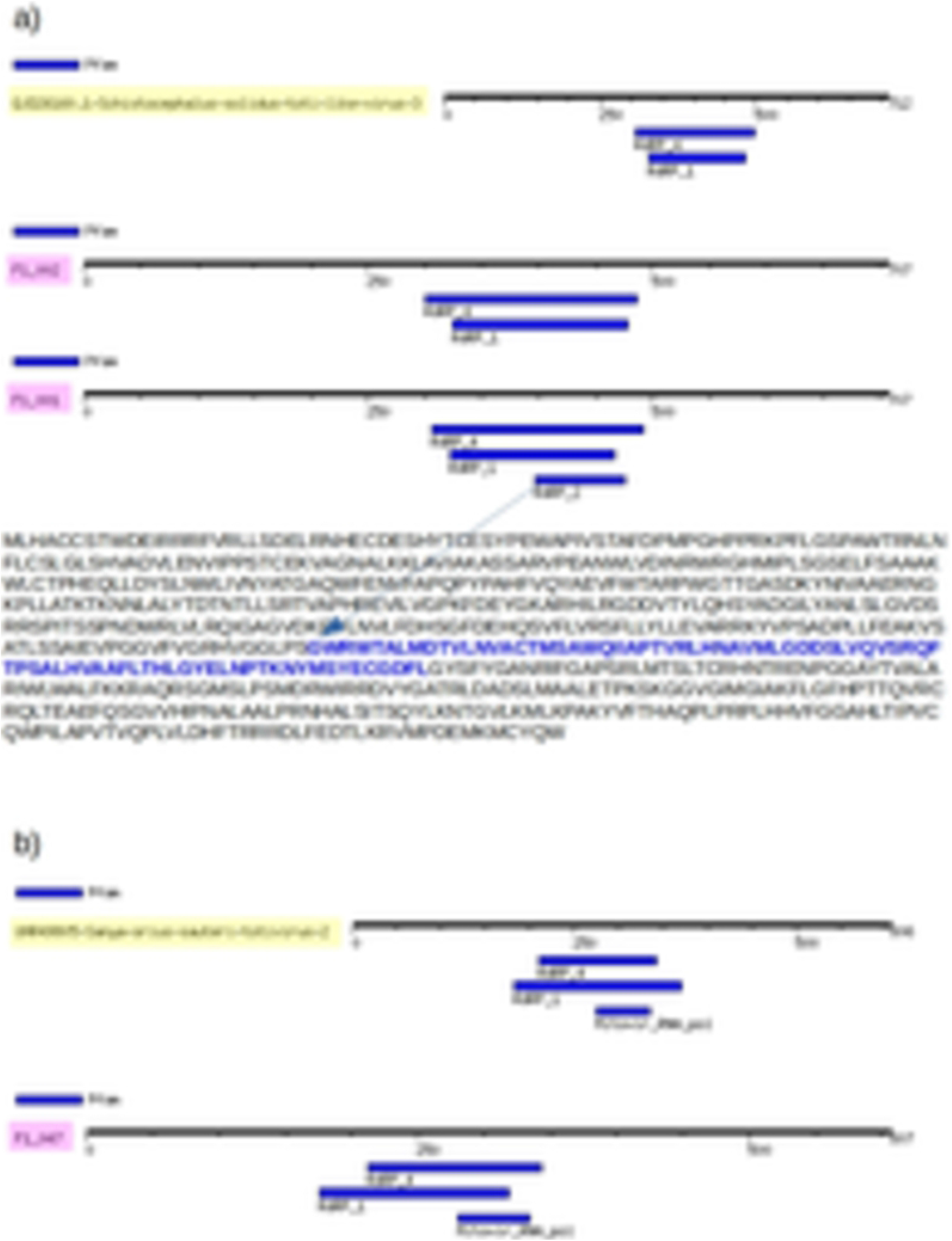
RdRpol motifs of totiviruses. The translated polymerase sequence of totiviruses is represented by horizontal bars that show the location of RdRpol motifs. The polymerase of F1_001 and F1_002 were compared to their best-hit reference QJD26160 (identified in Schistocephalus solidus), and the identified motifs are shown. In addition, F1_001 has an extra motif that is highlighted in blue and indicated by an arrow in this figure. In sequence F1_047, the motifs are compared to its cognate best-hit reference UHR49805, which was identified in the herbivorous insect *Eysarcoris guttigerus* that feeds on plants.

### Phylogenetic tree of RdRpol of totiviruses

Phylogenetic inference was performed using amino acid sequences of the RdRpol region to classify the sequences generated in this study. To this end, we utilized selected 53 reference sequences from the Totiviridae family. The resulting phylogenetic tree (Fig 3) was generated using a Coalescent Bayesian approach. The tree accurately depicts the major genera of totiviruses, including *Giardiavirus, Trichomonavirus, Leishmaniairus, Victorivirus*, and *Totivirus*. Furthermore, the recently proposed genera *Artivirus, Pistolvirus* and *Tricladivirus* are also identified in this tree. The sequences from this study were grouped into two distinct phylogenetic clades in the tree (indicated in blue color in the tree of Figure 3). Most sequences (i.e., F1_001, F1_006, F1_032, F1_066, F1_046, F1_052, F1_010, F1_012, F1_051, F1_042, F1_079, F1_045, F1_058, F1_043, F1_044, F1_036, F1_003, F1_062, F1_021, F1_007, F1_048, F1_081, F1_064, F1_008 and F1_002) clustered in a clade (here named

**Fig 3.**
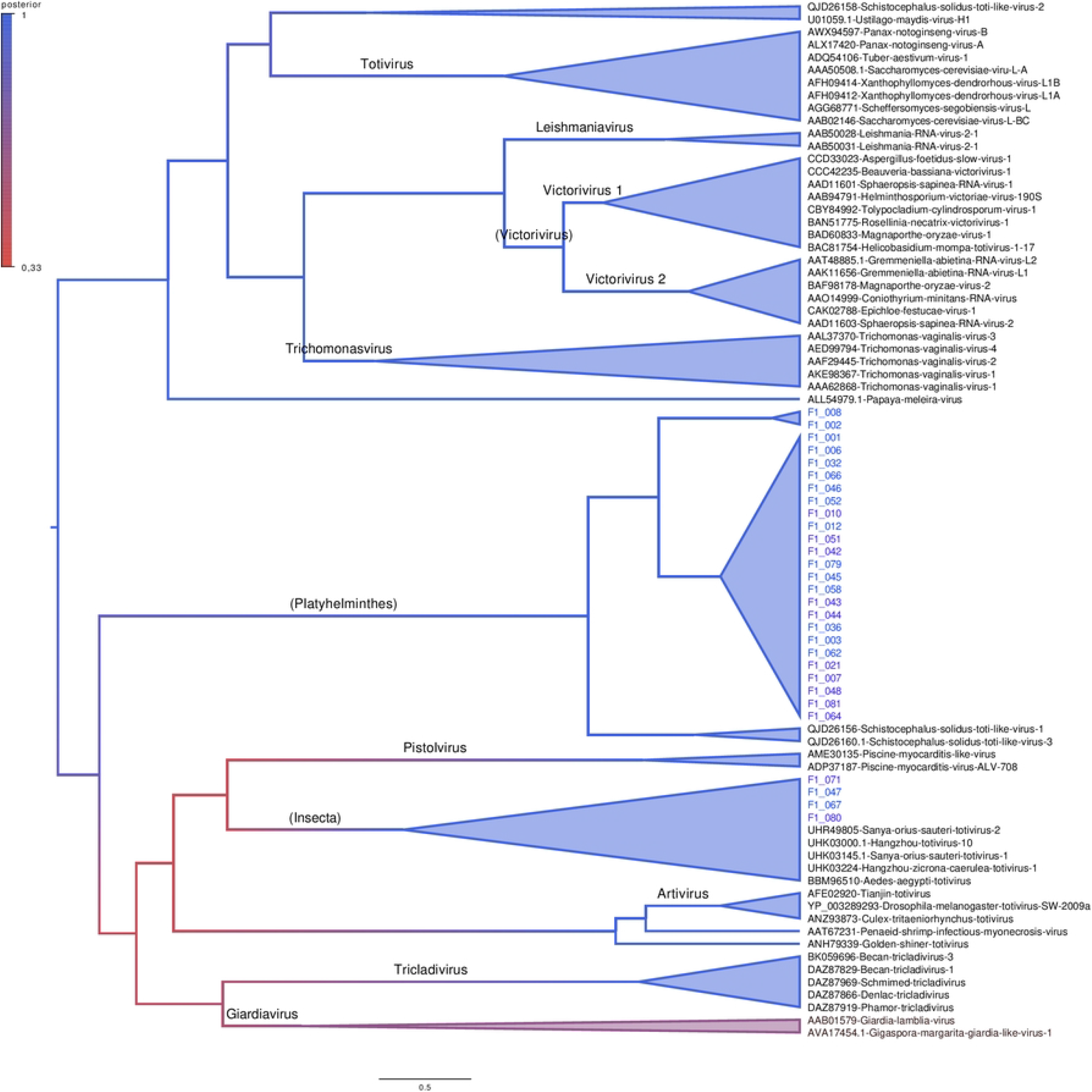
Phylogenetic tree of RdRpol of totiviruses. The translated RdRpol sequences were used to infer the tree, and branch support is based on posterior probability, which is indicated by colors according to the scale in the upper part of the figure. Clades with high branch support are represented by triangles in the tree. The horizontal bar shows the scale of the tree in substitutions per site. Sequences identified in this study are indicated in blue color in the tree. The names above the branches indicate the main totivirus genera as per the International Committee on Taxonomy of Viruses (ICTV). Platyhelminthes and Insecta are not genera; rather, they represent clades identified in this study.

Platyhelminthes) that includes the sequences QJD26156 and QJD26160 identified previously in the platyhelminthes *Schistocephalus solidu*s in the United States in 2018. The sequences F1_071, F1_047, F1_067 and F1_080 form a cluster (named Insecta) with the reference UHR49805 that was identified in the insect *Eysarcoris guttigerus* in China in 2017.

### RdRpol amino acid identity

Our phylogenetic analysis revealed that the sequences in the Platyhelminthes clade generated in this study could be categorized into two subclades. To further examine the level of identity of the RdRpol of these sequences, we created a subtree containing the Platyhelminthes clade and used it to illustrate the level of identity (Fig 4). Amino acid identity between F1_008 and its sister sequence F1_002 was found to be 98%. The mean identity among sequences in the clade composed of F1_001, F1_006, F1_032, F1_066, F1_046, F1_052, F1_010, F1_012, F1_051, F1_042, F1_079, F1_045, F1_058, F1_043, F1_044, F1_036, F1_003, F1_062, F1_021, F1_007, F1_048, F1_081, and F1_064 is 94%. In contrast, the identity between the sequences identified in *Schistocephalus solidus* is only 59%. On the other hand, the mean identity of sequences identified in *Schistocephalus solidus* with F1_001 and F1_002 is 42% and 41%, respectively. The identity between F1_001 and F2_002 is 54%.

**Fig 4.**
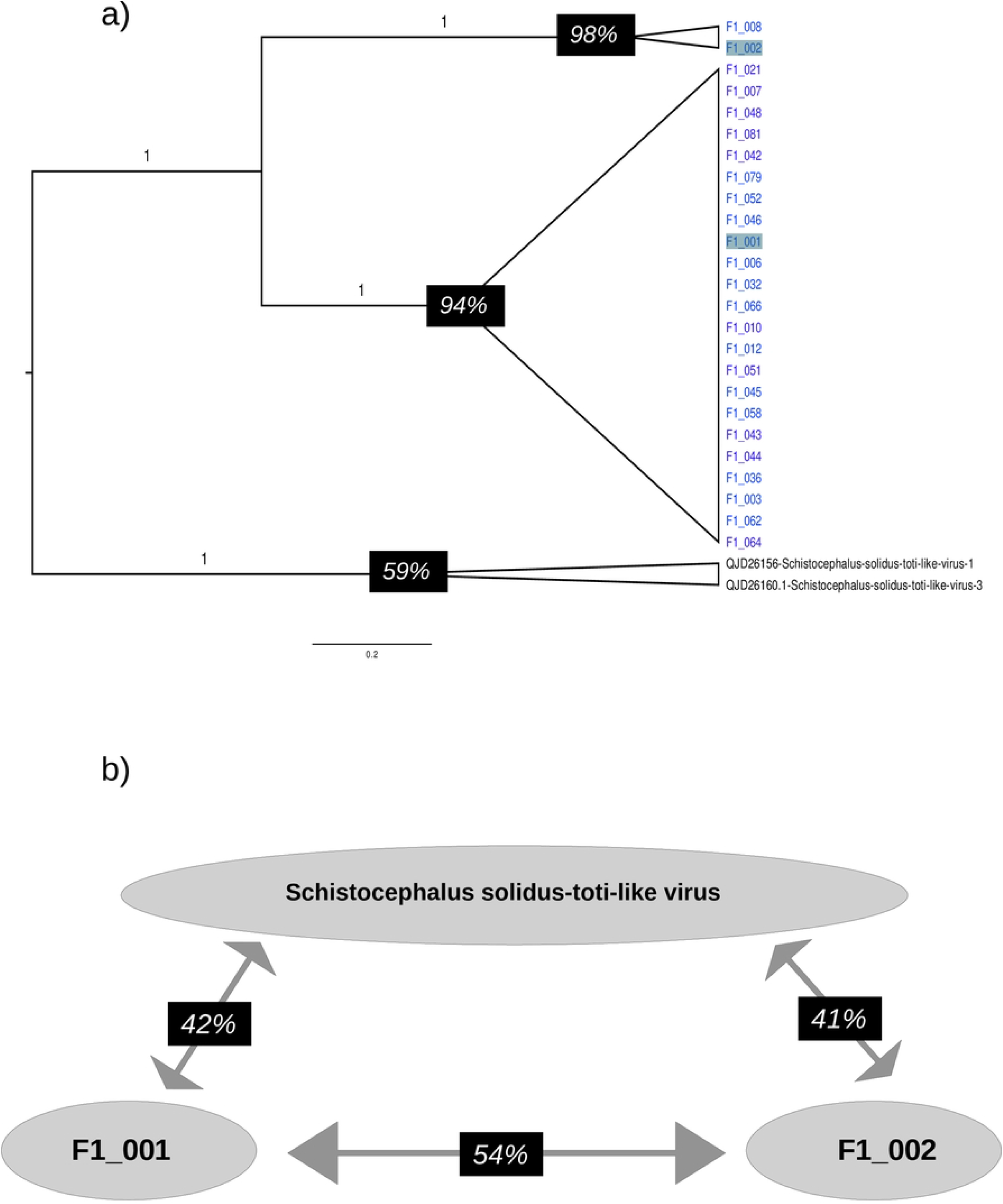
Amino acid identity in the RdRpol of totiviruses. a) Subtree depicting the phylogenetic clade Platyhelminthes. Values above branches indicate the posterior probability. The identity within each clade is indicated by black rectangles in the tree. b) Identity of sequences identified in *Schistocephalus solidus* with F1_001 and F1_002.

## Discussion

Viral surveillance in bats has been conducted extensively in southern Brazil, but few studies have explored bat viruses in other regions. Previous investigations in Brazil have identified Flavivirus, Coronavirus, Arenavirus, Paramyxovirus, Adenovirus, Papillomavirus and Parvovirus in the bats *Molossus molossus, Artibeus lituratus* and *Sturnira lilium* [7,9,14,14,16,20,28,29,63,64].

The emergence of next-generation sequencing technologies has allowed for the detection of many viruses in bats [6,25–27,27,65]. There are few reports of totiviruses in bats [27,32]. Recently, two contigs (2059bp and 627bp) of a totivirus-like virus were identified in the insectivorous bat *Nyctalus noctula* during a study on bat carcasses [31]. Similarly, short reads of totivirus were found in the feces of *Molossus molossus* bats in French Guiana in another study [30]. It is important to note that in metagenomics studies, it is not possible to determine the exact viral host due to the nature of next-generation sequencing approaches. This is also true for totiviruses identified in bats; their host can be ecto- or endoparasites, or even the food source consumed by these animals. The only totivirus-like virus that has been isolated in insect cells, named Tianjin totivirus, was detected in the guano of Myotis bats [32]. It is believed that this virus is likely to infect insects that are consumed by insectivorous bats.

We discovered 47 contigs in liver samples of Molossus bats that exhibit varying degrees of amino acid similarity with totivirus-like sequences previously detected in diverse hosts. Our thorough phylogenetic analysis revealed twenty-five RdRpol sequences that are closely linked to the platyhelminthes *Schistocephalus solidus* found in the United States in 2018. Additionally, we identified four more RdRpol sequences associated with a totivirus reference discovered in the insect *Eysarcoris guttigerus* in China in 2017. It is crucial to highlight that the amino acid identity between our sequences and the totivirus-like reference is below 50%, indicating significant genetic divergence between these viruses. Notably, the Tianjin totivirus, isolated from guano bats, exhibits less than 18% amino acid identity in the RdRpol region compared to the totivirus-like sequences identified in Molossus bats. This observation strongly suggests that these viruses are ancient and possess host-specific characteristics.

In this study, we conducted next-generation sequencing to identify totivirus-like viruses in liver tissue obtained from *Molossus molossus* bats residing in the Amazon region of Brazil. Through comparative phylogenetic analysis of the RNA-dependent RNA polymerase region, we discovered that the viruses found in Molossus bats belong to two distinct phylogenetic clades, potentially representing different genera within the Totiviridae family. These findings indicate that the diversity of totiviruses in bats surpasses previous estimations and underscore the significant role of bats as potential reservoirs for novel toti-like viruses.

## Author Contributions

Conceptualization, W.U.A, L.R.R.R, L.F.M, E.L and A.C.d.C.; methodology, R.d.S.C, E.d.S.F.R, W.U.A, L.R.R.R, L.F.M, V.d.S.M, F.V, X.D, E.D, E.L and A.C.d.C.; investigation, R.d.S.C, E.d.S.F.R, W.U.A, L.R.R.R, L.F.M, V.d.S.M, F.V, R.P.P, X.D, E.D, E.L and A.C.d.C.; data curation, R.d.S.C, E.d.S.F.R, W.U.A, L.R.R.R, L.F.M, V.d.S.M, F.V, R.P.P, X.D, E.D, E.L and A.C.C.; writing—original draft preparation, E.d.S.F.R, E.L.; writing— review and editing, R.P.P.; supervision, E.L and A.C.d.C X.X.

## Acknowledgments

We thank Coordenação Geral de Laboratórios de Saúde Pública do Departamento de Articulação Estratégica da Secretaria de Vigilância em Saúde do Ministério da Saúde (CGLAB/DAEVS/SVS-MS), MP Biomedicals do Brasil, Zymo Research Inc. for the donation of reagents for this project. We thank Luciano Monteiro da Silva and Nilton Costa. We thank the Pró-reitoria de pesquisa e pós-graduação of UFPA for supporting the publication costs.

